# Minimal effective dose of telacebec administered orally in a murine model of leprosy

**DOI:** 10.64898/2026.06.05.730402

**Authors:** Chauffour Aurélie, Avanzi Charlotte, Godmer Alexandre, Poignon Corentin, Natalya Serbina, Cambau Emmanuelle, Bettina E. Broeckling, Joseph Himelspach, John Anderson, Veziris Nicolas, Aubry Alexandra

## Abstract

Leprosy treatment requires prolonged therapy with challenging patient follow-up. New regimens are needed to simplify current treatments. A recent clinical trial evaluating bedaquiline has shown promising results; however, to prevent the emergence of drug resistance, additional therapeutic options are required. Telacebec (TCB), an imidazopyridine amide targeting the *Mycobacterium leprae* electron transport chain, represents a promising candidate. In this study, we determined the minimal effective dose (MED) of TCB against *M. leprae in vivo* by using the proportional bactericidal method in the mouse footpad model. Results were analyzed by using microscopy, RLEP qPCR and molecular viability. The MED obtained was 20 mg/kg. TCB doses ≥20 mg/kg achieved complete bacterial clearance, similar to bedaquiline 25mg/kg. Molecular enumeration confirmed these findings whereas molecular viability assessment had limited applicability due to insufficient bacterial burden. These findings provide a strong foundation for clinical trial design and the development of combination therapies.

## INTRODUCTION

Among mycobacterial diseases, leprosy is the second most prevalent after tuberculosis. The disease is caused by *Mycobacterium leprae* and *M. lepromatosis* [1]. A major challenge in leprosy research is that both pathogens cannot be cultivated *in vitro*, greatly complicating the evaluation and development of antileprosy drugs. The first effective treatment, dapsone, was introduced in the 1940s [2,3]. Subsequently, clofazimine and rifampicin (RIF) were shown to possess bactericidal activity against *M. leprae*, leading the World Health Organization (WHO) to recommend a multidrug therapy (MDT) combining these three agents [4–6]. Although highly effective, this regimen requires prolonged administration (12–24 months) and remains difficult to implement and monitor in endemic settings, underscoring the urgent need for shorter and more effective therapeutic strategies [7].

Advances in leprosy treatment have largely paralleled those achieved in tuberculosis. A notable example is bedaquiline (BDQ), which targets the c subunit of the F_0_F_1_ ATP synthase within the bacterial electron transport chain [8]. Owing to the markedly reduced electron transport chain of *M. leprae*, BDQ demonstrated exceptional activity in a murine model of leprosy, with a minimal effective dose of 3.3 mg/kg [7,8]. These preclinical findings were supported by two clinical trials confirming its efficacy against the leprosy bacillus [9].

Telacebec (TCB) or Q203, is a novel imidazopyridine amide compound also targeting the electron transport chain, and more precisely the cytochrome bc_1_-aa_3_ complex [10]. TCB has shown potent activity against *M. tuberculosis* and *M. ulcerans in vitro* and in a phase 2 clinical trial for *M. tuberculosis* [11–14]. Similarly to BDQ, TCB monotherapy (2mg/kg) was demonstrated to exhibit *ex vivo* and *in vivo* activity against *M. leprae* comparable to that of MDT, suggesting important potential for leprosy patient management and treatment duration reduction [15]. Building on these promising findings, we aimed to determine the minimal effective oral dose (MED) of TCB in a relevant mouse model of leprosy.

## RESULTS

### MED determined by the proportional bactericidal method

Eleven months after inoculation, all mice were positive in the untreated control groups within the inocula 5×10^4^ Acid Fast Bacilli per footpad (AFB/ footpad) until 5×10^2^ AFB/ footpad; 5 mice among 7 in the 5.10^1^ group were positive; whereas all mice were negative in the 5.10^0^ group (table 1 and figure 1). The proportion of viable *M. leprae* bacilli in the untreated group was 2.26% among the initial suspension inoculated (table 1).

**Table 1.**
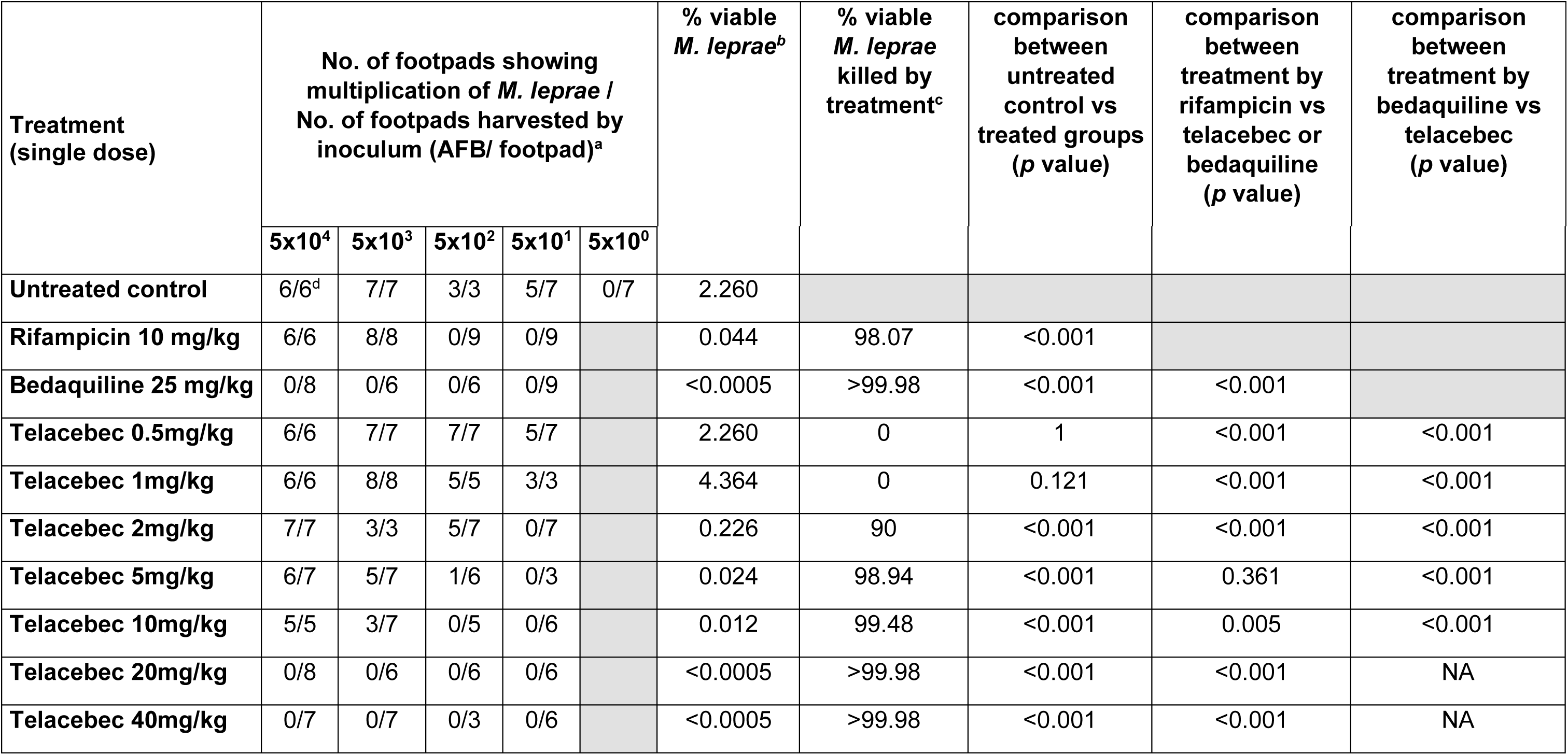
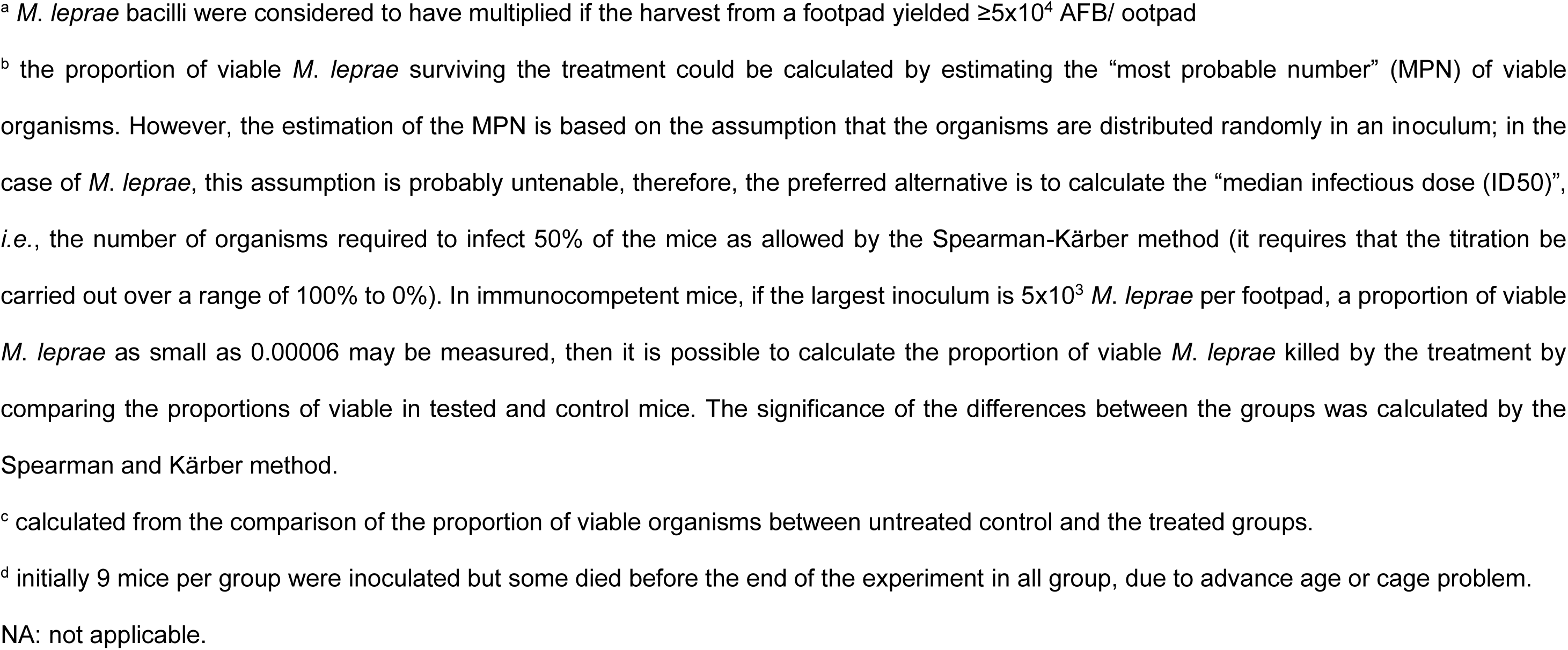
Bactericidal activity of a dose-ranging of telacebec against *M. leprae* THAI53 measured in nude mice by the proportional bactericidal method.

| Treatment<br>(single dose) | No. of footpads showing<br>multiplication of *M. leprae* /<br>No. of footpads harvested by<br>inoculum (AFB/ footpad)<sup>a</sup> |  |  |  |  | % viable<br>*M. leprae*<sup>b</sup> | % viable<br>*M. leprae*<br>killed by<br>treatment<sup>c</sup> | comparison<br>between<br>untreated<br>control vs<br>treated groups<br>(*p* value) | comparison<br>between<br>treatment by<br>rifampicin vs<br>telacebec or<br>bedaquiline<br>(*p* value) | comparison<br>between<br>treatment by<br>bedaquiline vs<br>telacebec<br>(*p* value) |
| --- | --- | --- | --- | --- | --- | --- | --- | --- | --- | --- |
|  | 5x10<sup>4</sup> | 5x10<sup>3</sup> | 5x10<sup>2</sup> | 5x10<sup>1</sup> | 5x10<sup>0</sup> |  |  |  |  |  |
| Untreated control | 6/6<sup>d</sup> | 7/7 | 3/3 | 5/7 | 0/7 | 2.260 |  |  |  |  |
| Rifampicin 10 mg/kg | 6/6 | 8/8 | 0/9 | 0/9 |  | 0.044 | 98.07 | <0.001 |  |  |
| Bedaquiline 25 mg/kg | 0/8 | 0/6 | 0/6 | 0/9 |  | <0.0005 | >99.98 | <0.001 | <0.001 |  |
| Telacebec 0.5mg/kg | 6/6 | 7/7 | 7/7 | 5/7 |  | 2.260 | 0 | 1 | <0.001 | <0.001 |
| Telacebec 1mg/kg | 6/6 | 8/8 | 5/5 | 3/3 |  | 4.364 | 0 | 0.121 | <0.001 | <0.001 |
| Telacebec 2mg/kg | 7/7 | 3/3 | 5/7 | 0/7 |  | 0.226 | 90 | <0.001 | <0.001 | <0.001 |
| Telacebec 5mg/kg | 6/7 | 5/7 | 1/6 | 0/3 |  | 0.024 | 98.94 | <0.001 | 0.361 | <0.001 |
| Telacebec 10mg/kg | 5/5 | 3/7 | 0/5 | 0/6 |  | 0.012 | 99.48 | <0.001 | 0.005 | <0.001 |
| Telacebec 20mg/kg | 0/8 | 0/6 | 0/6 | 0/6 |  | <0.0005 | >99.98 | <0.001 | <0.001 | NA |
| Telacebec 40mg/kg | 0/7 | 0/7 | 0/3 | 0/6 |  | <0.0005 | >99.98 | <0.001 | <0.001 | NA |

<sup>a</sup> *M. leprae* bacilli were considered to have multiplied if the harvest from a footpad yielded $\geq 5 \times 10^4$ AFB/ ootpad
<sup>b</sup> the proportion of viable *M. leprae* surviving the treatment could be calculated by estimating the “most probable number” (MPN) of viable organisms. However, the estimation of the MPN is based on the assumption that the organisms are distributed randomly in an inoculum; in the case of *M. leprae*, this assumption is probably untenable, therefore, the preferred alternative is to calculate the “median infectious dose (ID50)”, *i.e.*, the number of organisms required to infect 50% of the mice as allowed by the Spearman-Kärber method (it requires that the titration be carried out over a range of 100% to 0%). In immunocompetent mice, if the largest inoculum is $5 \times 10^3$ *M. leprae* per footpad, a proportion of viable *M. leprae* as small as 0.00006 may be measured, then it is possible to calculate the proportion of viable *M. leprae* killed by the treatment by comparing the proportions of viable in tested and control mice. The significance of the differences between the groups was calculated by the Spearman and Kärber method.
<sup>c</sup> calculated from the comparison of the proportion of viable organisms between untreated control and the treated groups.
<sup>d</sup> initially 9 mice per group were inoculated but some died before the end of the experiment in all group, due to advance age or cage problem.
NA: not applicable.

**Figure 1.**
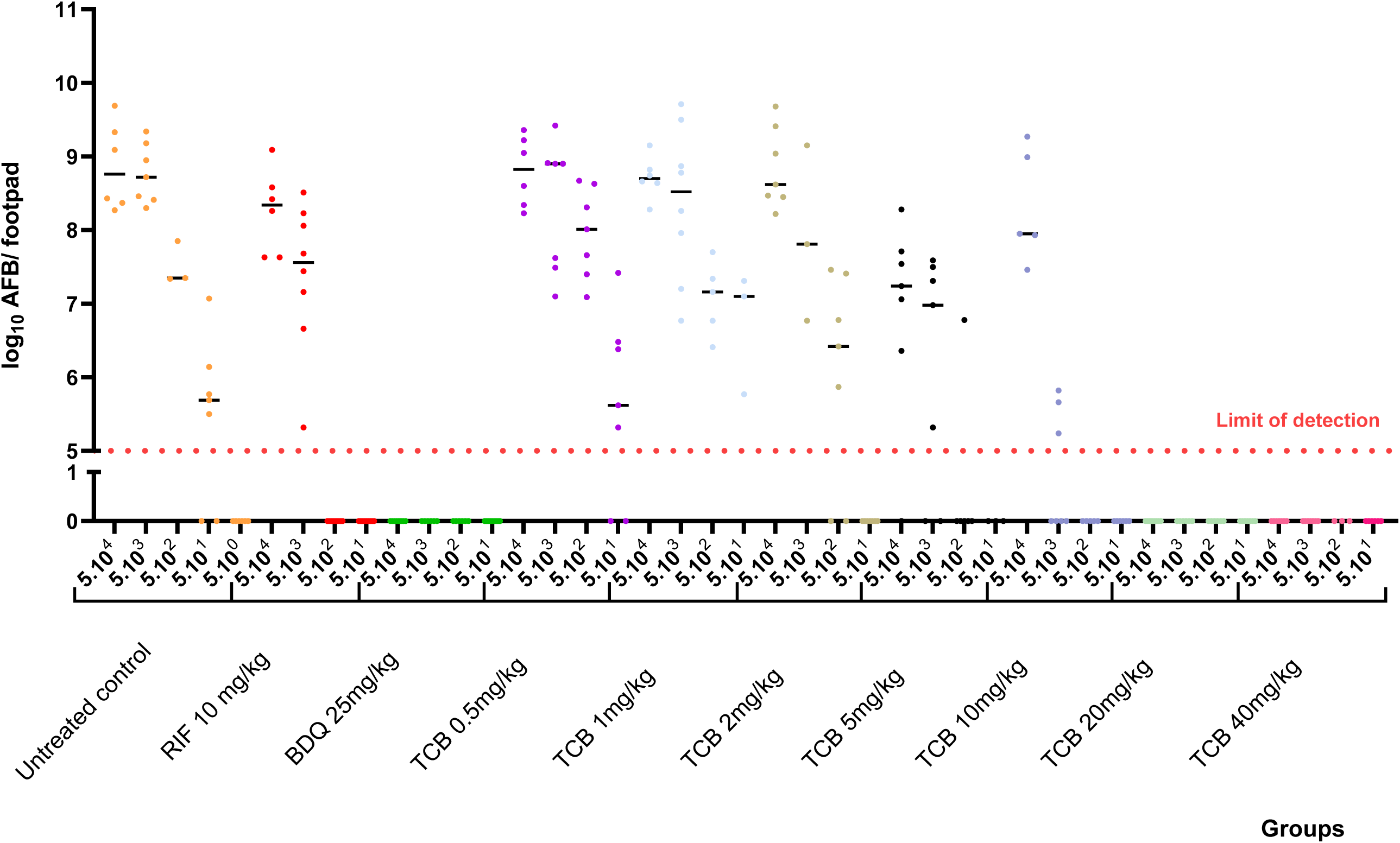
Bacterial load of *M. leprae* in nude mice treated with different doses of telacebec vs untreated control group. Each dot represents a mouse and the dotted red line indicates the threshold of detection of *M. leprae*.

In the RIF and BDQ groups, the percentages of viable bacilli were significantly lower than in the untreated control group with 0.044% and <0.0005% respectively (*p* < 0.001) (table 1). Regarding the TCB-treated groups, there was a ranking efficacy depending on the dose: the lowest doses, *i.e.*, 0.5 and 1mg/kg, demonstrated no activity against *M. leprae*, whereas bacilli started to be killed at 2mg/kg of TCB (0.226% of viable bacilli *vs.* 2.260% in the untreated control group; *p* <0.001; table 1). Among the doses enabling to kill bacilli, 10, 20 and 40mg/kg were more effective than RIF whereas 20 and 40mg/kg were as effective as BDQ (table 1, figure 1). Notably, these latter doses cleared AFB in all mouse footpads, as BDQ. The MED of telacebec is therefore 20mg/kg in our murine model of leprosy.

### MED determined by molecular enumeration

Bacterial enumeration was also performed using RLEP qPCR to quantify *M. leprae* burden, with Ct values converted to log₁₀ *M. leprae*/mL using established standard curves (Supplementary Data Table S1-3). The effect of each treatment on bacterial log_10_ *M. leprae*/mL after 1- and 11-months post-treatment was assessed relative to the untreated control at two inoculum concentrations (5×10^4^ and 5×10^3^ AFB/ footpad). No differences among groups were observed after 1 month. As normality assumptions were not consistently met, Kruskal-Wallis tests were used, revealing significant differences among groups (5×10^3^: H = 33.05, p < 0.001; 5×10^4^: H = 36.23, p < 0.001) (Supplementary Data Table 1).

Following the Kruskal-Wallis test, pairwise comparisons were limited to treatment *versus* control contrasts and performed using Mann-Whitney U tests with Benjamini-Hochberg FDR - adjusted *p-*values (Supplementary Data table S3 and figure 2). At the 5×10³ AFB/ footpad inoculum, the untreated control reached 9.47 ± 0.34 log_10_ *M. leprae*/mL. Low TCB concentrations (0.5–2mg/kg) and RIF 10mg/kg did not significantly reduce bacterial counts compared to the untreated group (all adjusted *p*-values > 0.05). TCB-5 and TCB-10 showed a numerical, though non-significant, reduction in bacterial burden (7.63 ± 2.29 and 8.10 ± 1.84 log_10_ *M. leprae*/mL, respectively; adjusted *p-*values = 0.21 and 0.73). At the 5×10⁴ AFB/ footpad inoculum (untreated control: 9.53 ± 0.34 log_10_ *M. leprae*/mL), the pattern of activity was fully consistent with that observed at the lower inoculum (Supplementary Data table S3, figure 2). The reproducibility of effect size and statistical significance across both inoculum concentrations indicates that the bactericidal activity of TCB ≥20 mg/kg and BDQ-25 is not contingent on the initial bacterial load. Under the conditions tested, 20 mg/kg represented the minimum efficacious concentration: TCB-20, TCB-40, and BDQ-25 were the only groups to achieve a statistically significant reduction, decreasing bacterial burden by approximately 5 log_10_ *M. leprae*/mL relative to the untreated control (3.89 ± 0.86, 4.13 ± 0.99, and 4.37 ± 0.65 log_10_ *M. leprae*/mL, respectively; adjusted *p-*values = 0.024 for each).

**Figure 2:**
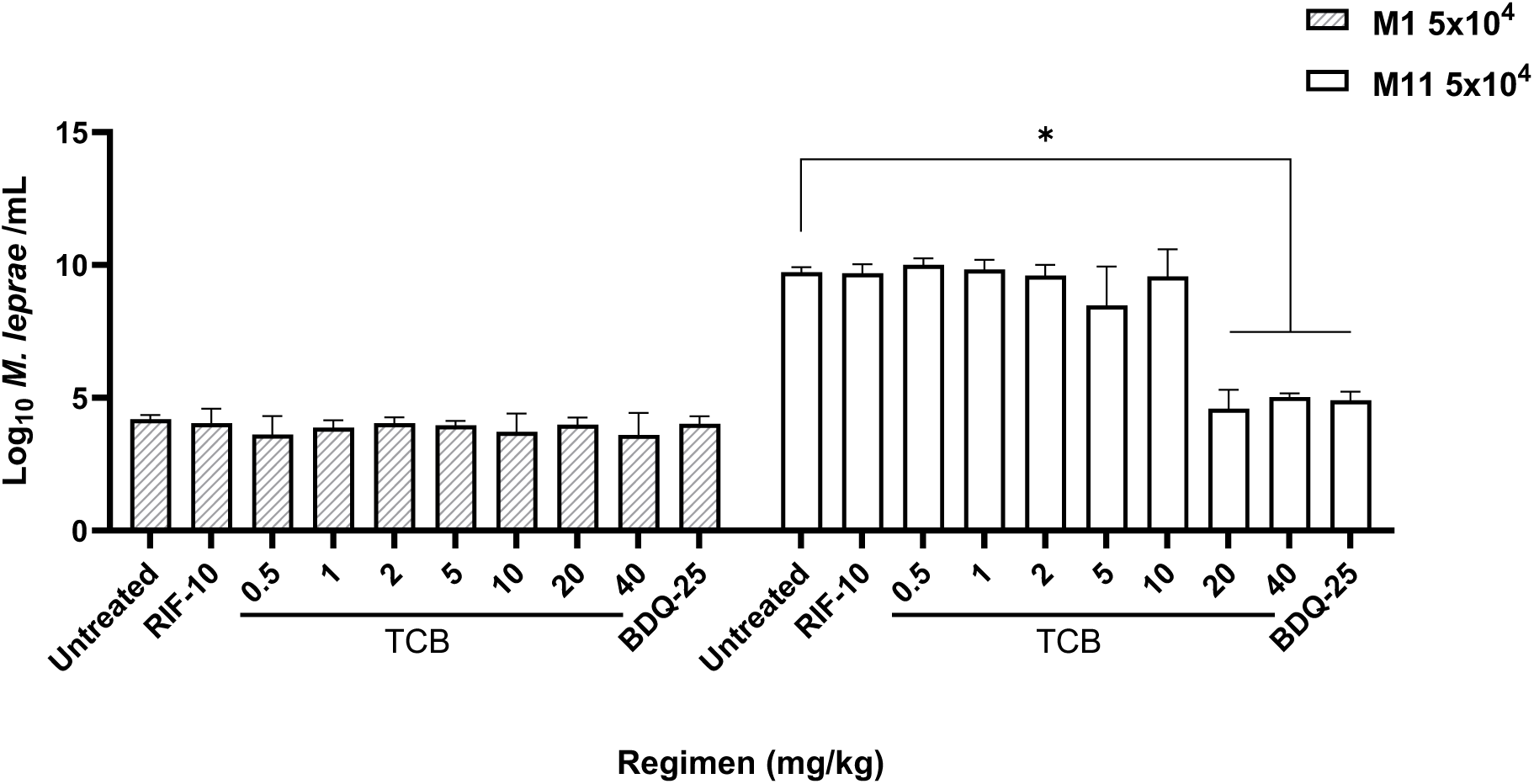
Reduction in bacterial burden following treatment in the mouse footpad model measured by RLEP qPCR. Bacterial burden (log_10_ bacilli / ml) was quantified following treatment across the inoculum 5×10⁴ AFB/ footpad over time (*i.e.,* 1- and 11-months after treatment). Horizontal bars indicate the median. Statistical analyses were performed separately for each period using one-way ANOVA on log_10_-transformed data, followed by multiple comparisons against the untreated control group. Adjusted *p-*values are indicated as follows: *p* < 0.05 (*), *p* < 0.01 (**), *p* < 0.001 (***), *p* < 0.0001 (****). High-dose treatments (TCB 20–40 mg/kg and BDQ) resulted in substantial reductions in bacterial burden, approaching 5–6 log_10_ decreases compared to untreated controls. All analyses were performed using GraphPad Prism v11.0.0. Analyses were also performed at month 11 on the 5×10^3^ groups and are presented in the supplementary data section.

### MED determined by molecular viability

Molecular viability assessment (MVA) via the established protocol targeting 16S rRNA and the transcripts *esxA* and *hsp18* was performed where applicable following Collins *et al.* [16]. Comparative analysis was only possible for groups with sufficient bacterial burden (untreated, RIF, TCB-0.5 through TCB-10 at month 11). The 16S rRNA showed consistent results among groups, as expected for a control marker (Supplementary Data table S4). Similarly, the mRNA viability markers (*esxA*, *hsp18*) did not show significant differences between the TCB 0.5-10 treatment groups (two-sided Mann–Whitney U tests with Bonferroni correction given the limited number of pairwise tests; all *adjusted p* > 0.05). For *esxA*, mean logarithmic values were similar across all TCB dose groups and RIF, ranging from 3.163 to 3.586 at 5×10^3^ and from 3.198 to 3.439 at 5×10^4^ AFB/ footpad, compared to untreated controls of 3.346 ± 0.307 and 3.568 ± 0.223, respectively. For *hsp18*, mean logarithmic values were also similar across all TCB dose groups and RIF, ranging from 3.130 to 3.340 at 5×10^3^ and from 2.900 to 3.437 at 5×10^4^ AFB/ footpad, compared to untreated controls of 3.293 ± 0.309 and 3.342 ± 0.296, respectively.

Critically, MVA could not be performed at month 1 for all the groups and at month 11 for the most efficacious treatments (TCB-20, TCB-40, and BDQ) due to insufficient bacterial burden. This technical limitation meant that the viability of *M. leprae* after treatments with the greatest bactericidal activity by enumeration could not be precisely evaluated in molecular viability. Low bacterial burden in these highly active treatment groups is itself indirect evidence of profound bacterial killing but precluded definitive viability assessment where it would be most clinically relevant.

## DISCUSSION

New drug discovery for leprosy remains challenging, with most candidates emerging from tuberculosis rather than dedicated leprosy research pipelines. The discovery of compounds targeting the electron transport chain in *M. tuberculosis* has opened new therapeutic avenues for leprosy treatment [9,17]. This approach is particularly promising given that *M. leprae* possesses an extensively downsized electron transport chain compared to other mycobacteria, containing only the cytochrome bc1-aa3 complex without the cytochrome bd oxidase pathway [18]. This limited respiratory repertoire potentially increases *M. leprae*’s vulnerability to electron transport chain inhibition, highlighting this pathway as an attractive therapeutic target. Previous studies have demonstrated that BDQ, which targets ATP synthase, exhibits potent anti-leprosy activity and is currently in phase III clinical evaluation [19], [9]. In parallel, TCB, targeting the cytochrome bc1-aa3 complex, has emerged as another promising inhibitor of the electron transport chain. Preliminary *ex vivo* and *in vitro* studies using 2mg/kg TCB confirmed its potential against *M. leprae* [15]. Given the scarcity of leprosy clinical trials, we aimed to determine the minimal effective dose of TCB to inform future trial design and compare its efficacy with RIF, the current backbone of leprosy treatment, and BDQ, the most promising new candidate.

To achieve this objective, we employed the proportional bactericidal mice model, specifically designed to evaluate the bactericidal power of antimicrobial agents *against M. leprae*. Using this model, we found that TCB at 20 and 40 mg/kg demonstrated bactericidal activity similar to BDQ 25mg/kg (table 1, figure 1). The lowest tested doses (0.5 and 1mg/kg) showed no activity against *M. leprae*, with viable bacilli counts equivalent to untreated controls. Bactericidal activity began at 2 mg/kg, though RIF retained superior activity. TCB activity gradually increased from 5 mg/kg onwards, reaching maximum effectiveness at 20mg/kg, where it exceeded RIF’s bactericidal activity and matched BDQ’s performance.

Molecular enumeration findings corroborated the mouse footpad model results, confirming the TCB 20mg/kg as the minimum efficacious dose for statistically significant bacterial reduction (figure 2). Both methodologies demonstrated equivalent bactericidal activity between TCB-20/40 and BDQ 25, with all achieving ∼5 log_10_ reductions in bacterial burden.

The molecular viability assay using the MVA method demonstrated significant limitations in the proportional bactericidal murine model for leprosy. Given the relatively low starting inocula (5×10³ and 5×10⁴ AFB/ footpad), highly bactericidal treatments resulted in insufficient residual bacterial burden for adequate RNA normalization (3000 bacilli needed) after 1 month, and even 11 months (table S1). This model-dependent constraint meant that MVA could not be performed precisely where it would be most clinically relevant, in evaluating the most efficacious treatments (TCB 20, TCB 40, BDQ 25). The inherent design of this experimental system, which uses various initial bacterial loads to assess bactericidal activity, creates a paradox where successful treatments eliminate too many bacteria for molecular viability assessment. Combined with low mRNA signal sensitivity, these findings highlight that while MVA may have utility in evaluating active compounds in other models, their practical application in bactericidal efficacy models is fundamentally limited by the inverse relationship between treatment success and residual bacterial availability for analysis.

The prospect of combining BDQ and TCB, both targeting the electron transport chain at different sites, offers hope for improved leprosy treatment. However, potential antagonism must be considered, as both compounds inhibit the same overall pathway. Studies in *M. tuberculosis* have demonstrated antagonism between QcrB-targeting compounds and BDQ under certain conditions [20–22]. Additionally, a recent murine leprosy study suggested possible antagonism between TCB and clofazimine, potentially due to competitive mechanisms or pharmacokinetic interactions [15]. Therefore, despite no antagonism has been shown in *M. leprae* between BDQ and TCB, further *in vivo* studies should evaluate combinations of electron transport chain-targeting drugs to determine whether this pathway’s therapeutic promise can be fully realized in leprosy treatment.

In conclusion, this study establishes a comprehensive framework for TCB dose evaluation in leprosy, defining the complete dose-response spectrum from inactive (0.5-1mg/kg) through emerging activity (2-10mg/kg) to maximal bactericidal effect (≥20mg/kg). This dosing framework directly enables rational clinical trial design and supports the broader effort to develop shorter, more effective leprosy treatment regimens aligned with WHO elimination goals.

## METHODS

### Ethics

The experimental project was favorably evaluated by the ethics committee n°5 Charles Darwin localized at the Pitié-Salpêtrière Hospital (Paris, France). Clearance was given by the French Ministry of Higher Education and Research under the number APAFIS#30674-202103181532328 v6. Our animal facility received the authorization to carry out animal experiments (license number D75-13-08). The persons who carried out the animal experiments had followed a specific training recognized by the French Ministry of Higher Education and Research and follow the European and the French recommendations on the continuous training. The design of the experimental project followed the guidelines ARRIVE [23].

### MED determined by the proportional bactericidal method

406 four-weeks-old female nude NMRI mice (Rj:NMRI-*Foxn1^nu/nu^*) were purchased at Janvier Labs, Le Genest Saint Isle, France. Mice were inoculated in both hind footpads with 30µl of the *M. leprae* strain THAI53 according to Shepard and Mac Rae method [24] (Figure 3). This strain is fully susceptible to the common antileprosy drugs rifampicin, dapsone, clofazimine and fluoroquinolones [25]. The suspension used to prepare the inoculum was freshly prepared as previously described from a mouse already infected with the strain [19].

**Figure 3.**
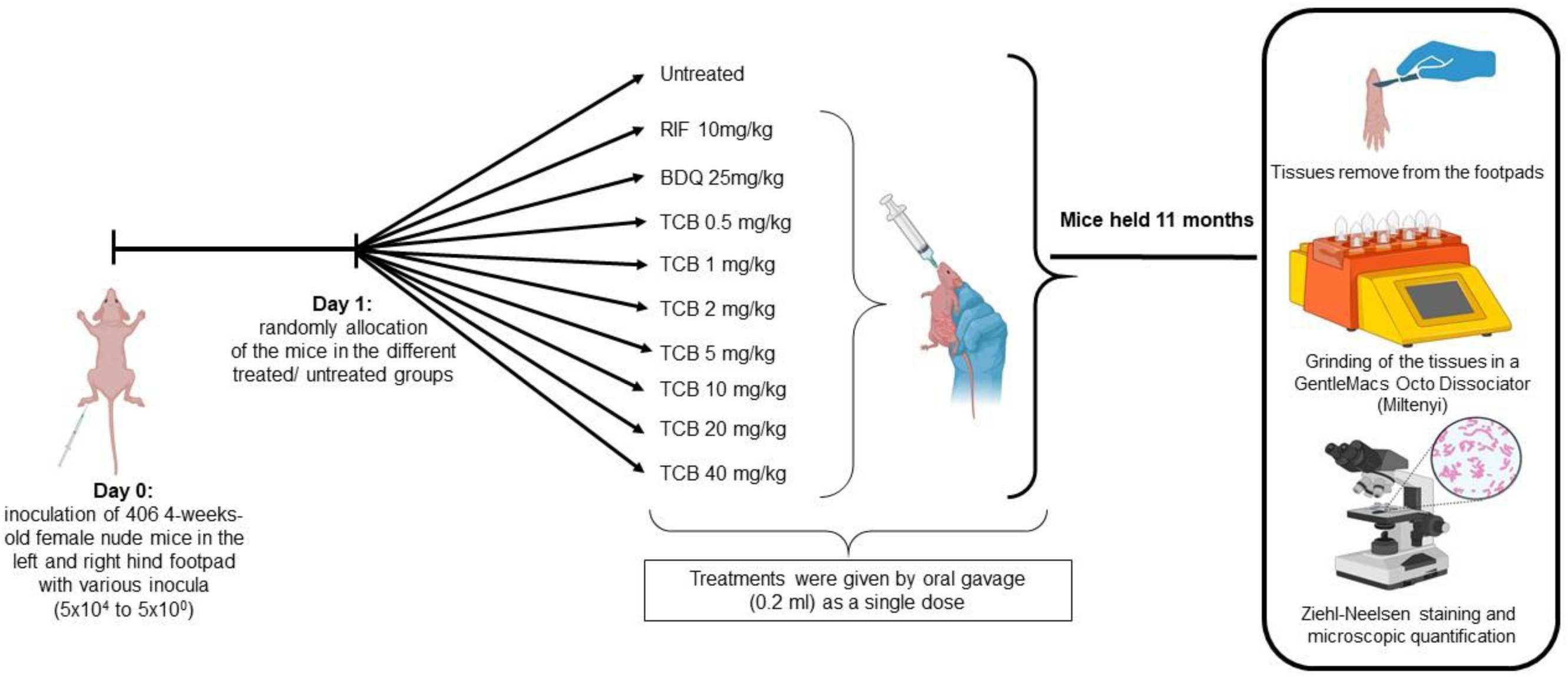
Protocol design of the in vivo MED of TCB study. Biorender credit (see license Created in BioRender. Aubry, A. (2026) https://BioRender.com/06oz8ar)

In this experiment, we used the proportional bactericidal method that allows measure the bactericidal activity of a compound. Mice were therefore inoculated with four different inocula of 5×10^4^, 5×10^3^, 5×10^2^, and 5×10^1^ AFB/ footpad, except for the untreated control group, which was also inoculated with a 5×10^0^ additional group.

After inoculation, mice were randomly allocated into one untreated control group and 9 treated groups of 9 mice each, except in the TCB 10, 20 and 40 mg/kg 5.10^1^ AFB/ footpads groups, where only 8 mice were inoculated, due to the death of 3 mice during the transport from the supplier to our animal facility. Treated groups were as follows: RIF 10 mg/kg, BDQ 25mg/kg, TCB within a dilution range from 0.5 to 40mg/kg. RIF is used as a control antibiotic representing the current standard anti-leprosy treatment, whereas BDQ serves as a control for the most promising new anti-leprosy agent identified to date. The day after inoculation, treatments were administered orally by gavage at a volume of 0.2 ml per mouse. Mice received a single dose for each treatment. RIF was purchased at Merck (France), whereas BDQ and TCB were kindly provided by the TB Alliance.

To permit multiplication of *M. leprae* to a detectable level, mice were held 11 months in the animal facility before euthanasia. The experiment usually lasts 12 months, but due to spontaneous death caused by the advanced age of the mice, we decided to shorten the experiment by one month. Tissues from mice footpad were then removed aseptically and homogenized under a volume of 2 ml of Hank’s balanced salt according to the Shepard’s method [26]. *M. leprae* bacilli were considered to have multiplied (*i.e.,* survived to the treatment) if those footpads were found to contain ≥5×10^4^ AFB, regardless of the size of the inoculum.

The proportion of viable *M. leprae* after treatment was determined from the infectious dose required to show multiplication in 50% of the inoculated mice. The significance of the differences between the groups was calculated by the Spearman and Kärber method [27]. A *p* <0.05 was considered statistically significant by standard evaluation. For multiple comparisons between the groups, Bonferroni’s correction was applied, *i.e.*, the difference would be significant at the 0.05 level only if the adjusted *p-* value adjusted to the number of groups: 0.05/n in which n was defined as the number of primary comparisons. Thus, the corrected *p* was 0.05/10 = 0.005 for the analysis comparing the treated groups to the untreated group and 0.05/8 = 0.0055 for the analysis comparing the RIF- or BDQ-treated groups to the other treated groups.

### MED determined by molecular enumeration and molecular viability

For molecular analyses, the contralateral footpad from the 5×10^4^ and 5×10^3^ groups was collected in DNA/RNA Shield (Zymo Research) at the time of euthanasia after 1 and/ or 11 months post-treatment. Each group comprised n = 5 independent replicates, except TCB-2 (n = 3 in 5×10^3^) and TCB-5 (n = 4 in 5×10^3^). After collection, tissues were mechanically disrupted using the gentleMACS™ Octo Dissociator system (Miltenyi Biotec) and stored at −80 °C prior to DNA and RNA extractions. These samples were subsequently used for molecular enumeration and viability assessment using the molecular viability assay (MVA).

RNA and DNA were extracted for all mouse footpad samples, and nucleic acid quality was assessed using an adapted protocol from Collins *et al.*, available on protocols.io (DOI: https://dx.doi.org/10.17504/protocols.io.q26g7ojmkvwz/v1) [16]. For each extraction batch, one uninfected mouse footpad was added to track cross-contamination. DNA was used for bacterial enumeration by RLEP qPCR as previously described [16]. DNase-treated RNA was used to quantify 16S rRNA, *esxA*, and *hsp18* transcripts using established protocols. For each target, standard curves were generated as described in previously, and all reactions were performed in technical duplicates. Negative controls included nuclease-free water, and positive controls consisted of plasmids as described previously [16]. RT-qPCR was performed on a CFX system (Bio-Rad), and data were analyzed using CFX Manager software.

Statistical analyses were performed separately for each inoculum using Prism v11.0.0. Prior to group comparisons, the normality of each treatment group was assessed using the Shapiro–Wilk test, and homogeneity of variances was evaluated with Levene’s test. As normality assumptions were not consistently met within each inoculum, the non-parametric Kruskal-Wallis test was used the non-parametric Kruskal-Wallis test was used to assess global differences among groups for each inoculum. Pairwise treatment *versus* control comparisons were subsequently performed using two-sided Mann-Whitney U tests. To control multiple comparisons across the nine treatment-*vs*.-control contrasts per inoculum, *p*-values were adjusted using the Benjamini-Hochberg false discovery rate (FDR) procedure. A significance threshold of α=0.05 was applied to the FDR-adjusted *p-*values. Results are reported as mean ± standard deviation. For molecular viability analyses, 16S rRNA levels were compared across treatment groups within each inoculum using a Kruskal–Wallis test, followed by pairwise two-sided Mann-Whitney U tests with Bonferroni correction for multiple comparisons (21 pairs). For *esxA* and *hsp18*, expression levels in each treatment group were compared to the untreated control using two-sided Mann-Whitney U tests with Bonferroni correction for multiple comparisons (n = 6 per inoculum). Statistical significance is indicated as: * *p* < 0.05, ** *p* < 0.01, *** *p* < 0.001.

## SUPPLEMENTARY DATA

**Table S1.** Pairwise comparisons of each treatment vs. untreated control by using RLEP qPCR at month 1. (Mann–Whitney U test, Benjamini–Hochberg FDR correction). Values are mean ± SD of log_10_ *M. leprae*/mL.

| Treatment | Inoculum 5×10<sup>4</sup> (control) |  |  |  |
| --- | --- | --- | --- | --- |
| | Mean ± SD | p (raw) | adjusted p-values | Sig. |
| Untreated | 4.2 ± 0.15 | | | |
| RIF-10 | 4.04 ± 0.55 | 0.6905 | 0.6905 | ns |
| TCB-0.5 | 3.62 ± 0.70 | 0.0159 | 0.1429 | ns |
| TCB-1 | 3.88 ± 0.29 | 0.0952 | 0.2857 | ns |
| TCB-2 | 4.05 ± 0.22 | 0.2222 | 0.3333 | ns |
| TCB-5 | 3.97 ± 0.17 | 0.0749 | 0.2857 | ns |
| TCB-10 | 3.72 ± 0.69 | 0.3095 | 0.3980 | ns |
| TCB-20 | 4.00 ± 0.26 | 0.2222 | 0.3333 | ns |
| TCB-40 | 3.60 ± 0.83 | 0.4206 | 0.4732 | ns |
| BDQ-25 | 4.02 ± 0.29 | 0.2222 | 0.3333 | ns |

\* p < 0.05 after FDR correction. ns: not significant.

**Table S2.** Shapiro–Wilk normality test results for enumeration by RLEP qPCR per treatment groups and inocula concentrations at month 11.

| Treatment | Inoculum 5×10<sup>4</sup> |  | Inoculum 5×10<sup>3</sup> |  |
| --- | --- | --- | --- | --- |
|  | Shapiro-Wilk p | Normal? | Shapiro-Wilk p | Normal? |
| Untreated | 0.4236 | yes | 0.5534 | yes |
| RIF-10 | 0.4190 | yes | 0.2224 | yes |
| TCB-0.5 | 0.7531 | yes | 0.6629 | yes |
| TCB-1 | 0.3028 | yes | 0.0612 | yes |
| TCB-2 | 0.5637 | yes | 0.8359 | yes |
| TCB-5 | 0.3165 | yes | 0.2654 | yes |
| TCB-10 | 0.2728 | yes | 0.0387 | NO |
| TCB-20 | 0.2469 | yes | 0.8571 | yes |
| TCB-40 | 0.0389 | no | 0.3967 | yes |
| BDQ-25 | 0.5925 | yes | 0.3850 | yes |

**Table S3.**
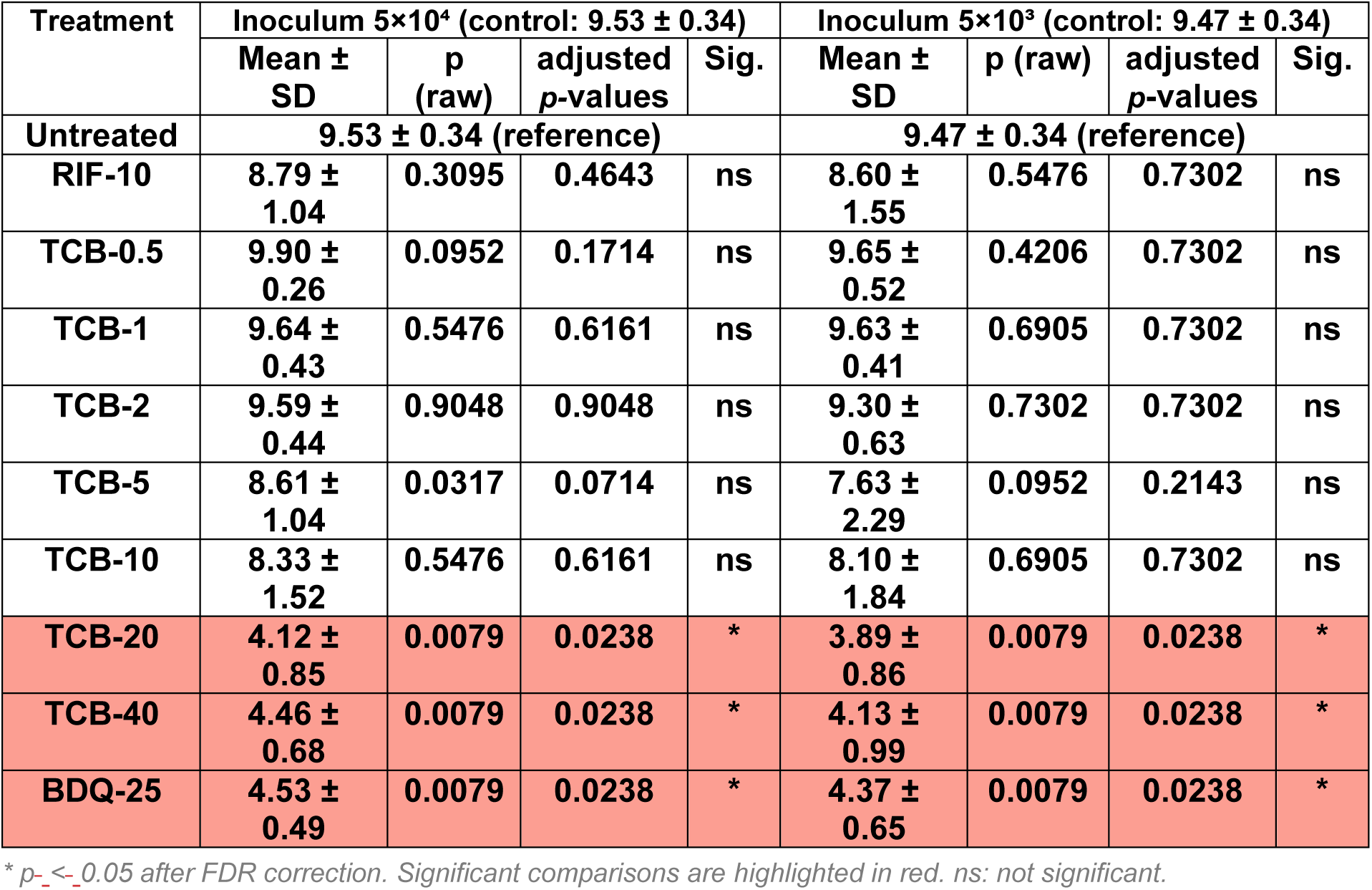
Pairwise comparisons of each treatment vs. untreated control by using RLEP qPCR at months 11. (Mann–Whitney U test, Benjamini–Hochberg FDR correction). Values are mean ± SD of log_10_ *M. leprae*/mL.

**Figure 4.**
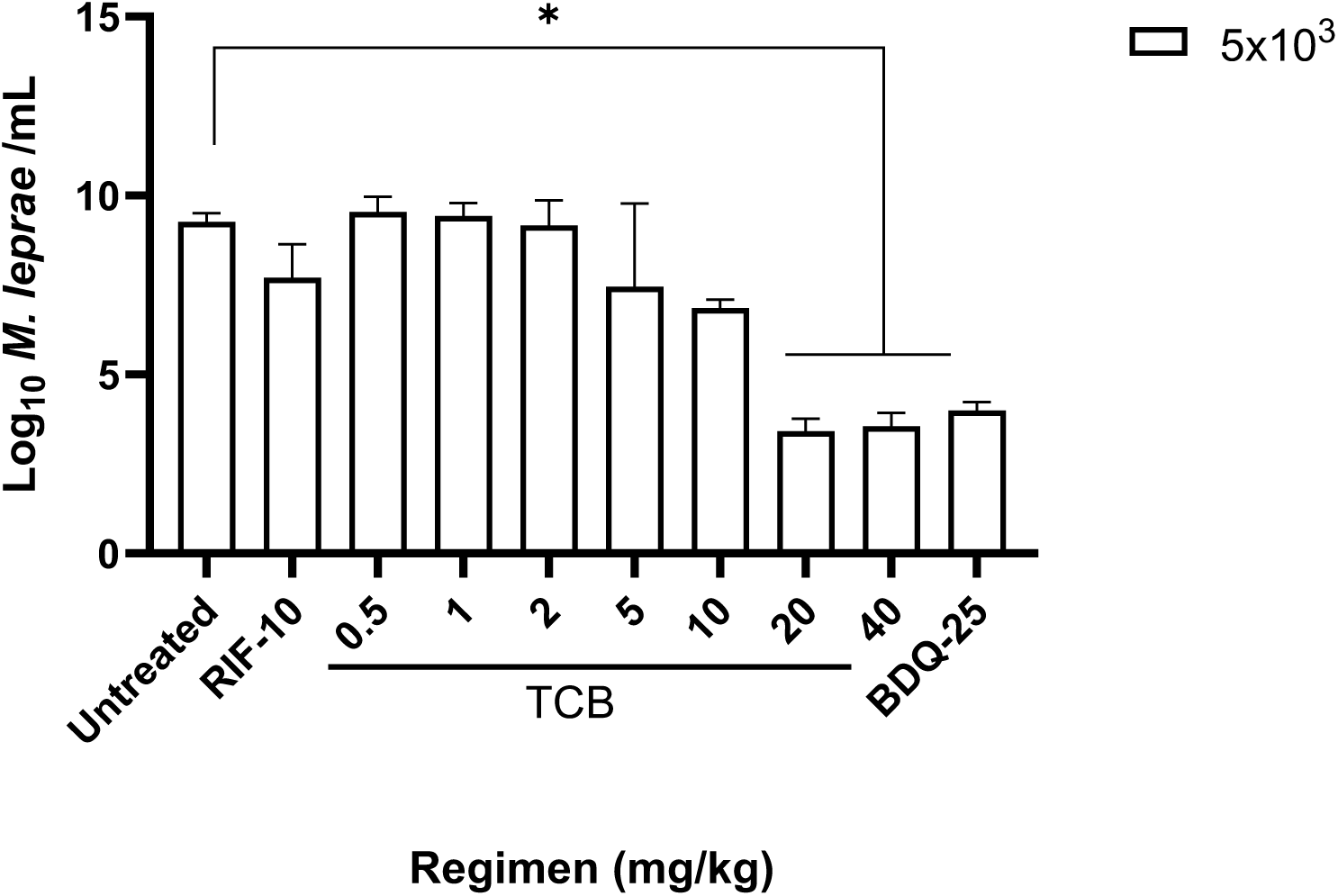
Enumeration of *M. leprae* by qPCR RLEP at month 11 in the 5×10^3^ groups.

Reduction in bacterial burden following treatment in the mouse footpad model measured by RLEP qPCR. Bacterial burden (log_10_ bacilli /mL) was quantified following treatment across the inoculum 5×10³ AFB/ footpad groups after 11 months of treatment. Horizontal bars indicate the median. Statistical analyses were performed separately for each inoculum period using one-way ANOVA on log₁₀-transformed data, followed by multiple comparisons against the untreated control group. Adjusted *p*-values are indicated as follows: *p* < 0.05 (*), *p* < 0.01 (**), *p* < 0.001 (***), *p* < 0.0001 (****). High-dose treatments (TCB 20–40 mg/kg and BDQ) resulted in substantial reductions in bacterial burden, approaching 5–6 log₁₀ decreases compared to untreated controls. All analyses were performed using GraphPad Prism v11.0.0.

**Table S4.** 16S rRNA copy numbers (mean ± SD) per treatment group and inoculum. Kruskal-Wallis omnibus test p-values and pairwise Bonferroni-corrected results are shown. The Kruskal-Wallis test revealed statistically significant overall variation across groups at both inocula (5×10^3^: H = 17.04, *p* = 0.009; 5×10^4^: H = 14.37, *p* = 0.026). However, after Bonferroni correction for 21 pairwise comparisons, no individual pair of groups reached significance at either inoculum.

| Treatment | n | 5×10<sup>4</sup> AFB/ footpad mean<br>Log<sub>10</sub> 16S ± SD | n | 5×10<sup>3</sup> AFB/<br>footpad mean<br>Log<sub>10</sub> 16S ± SD |
| --- | --- | --- | --- | --- |
| TCB-0.5 | 5 | 6.589 ± 0.192 | 5 | 6.638 ± 0.274 |
| TCB-1 | 5 | 6.565 ± 0.159 | 5 | 6.853 ± 0.195 |
| TCB-2 | 3 | 6.705 ± 0.102 | 5 | 6.887 ± 0.113 |
| TCB-5 | 3 | 6.500 ± 0.514 | 5 | 7.033 ± 0.038 |
| TCB-10 | 5 | 6.720 ± 0.145 | 5 | 6.284 ± 0.169 |
| RIF | 5 | 6.679 ± 0.158 | 5 | 6.761 ± 0.124 |
| Untreated | 5 | 7.059 ± 0.162 | 5 | 6.883 ± 0.247 |
| Kruskal-Wallis p |  | *p* = 0.026 |  | *p* = 0.009 |
| Pairwise (Bonferroni) |  | All ns |  | All ns |

*ns* = not significant after Bonferroni correction.

## Acknowledgments

We thank the Raoul Follereau Foundation for their financial support. We also thank the UMS 28 for their technical support.

## Author contributions

Funding acquisition: AA, CA. Conceived and designed the experiments: AA, AC, EC, AG, CP, NV. Performed the mouse experiments: AC, NV. Performed the molecular experiments: CA, BB, JH, JA. Supervision: AA. Analyzed the data: AA, AC, AG, BB, CA, CP, NV. Contributed reagents/materials/analysis tools: AC, CA, BB. Wrote the original draft: AA, AC, CA. Reviewed and edited the paper: all authors.

